# Polycystin-2 is cardioprotective against myocardial infarction by regulating the calcium-mediated ER stress response

**DOI:** 10.1101/2025.08.05.668747

**Authors:** Virdjinija Vuchkovska, Sanjeet Paluru, Ryne M. Knutila, Djamilla Simoens, Karla M. Márquez Nogueras, Elisabeth DiNello, Quan Cao, Jonathan A. Kirk, Ivana Y. Kuo

## Abstract

**Background:** Patients with autosomal polycystic kidney disease (ADPKD) have an increased risk and worsened outcomes for acute myocardial infarction (AMI), but the mechanism behind this is unknown. Polycystin 2 (PC2), the protein encoded by one of the two main genes mutated in ADPKD, is a calcium-permeant channel ubiquitously expressed; however, its role in cardiomyocytes remains poorly understood. One hallmark of AMI is ER stress, which PC2 is known to regulate adaptively; however, whether PC2 regulates ER stress in the ischemic heart is unknown.

**Objective:** This study investigates the mechanism by which PC2 regulates ER stress in myocardial ischemia.

**Methods and Results:** PC2 protein was increased in human ischemic heart failure samples and murine myocardial infarct samples, and it was enriched at ER-mitochondrial contact sites. Induction of myocardial infarction (MI) in cardiomyocyte-specific PC2-KO mice led to cardiac dysfunction and reduced PERK expression compared to control MI mice. ER stress induced by tunicamycin in vitro blunted PERK phosphorylation and subsequent CHOP upregulation in PC2 KO cells. Tunicamycin-induced ER stress resulted in a PC2-dependent ER calcium leak and mitochondrial calcium transients, along with increased mitochondrial function, all of which were decreased in PC2 KO cells. Moreover, PC2 KO cells after ER stress exhibited decreased mitochondrial membrane potential and increased apoptosis. Isolated WT cardiomyocytes exhibited increased diastolic calcium after acute ER stress induction and increased mitochondrial uptake, neither of which was seen in PC2 KO cells. Re-expression of full-length PC2 in vitro restored both the calcium leak and PERK phosphorylation in PC2 KO cells under ER stress, but not a pathological mutant PC2 D511V, which impairs ion channel activity.

**Conclusions:** PC2 is upregulated during ER stress, where it localizes at ER-mitochondrial contact sites and acts as an ER calcium leak channel, thereby restoring cellular homeostasis during the adaptive phase of ER stress. PC2 provides cardioprotection during ischemic events by preventing maladaptive ER stress, which contributes to cardiac dysfunction.

**Graphical abstract:** 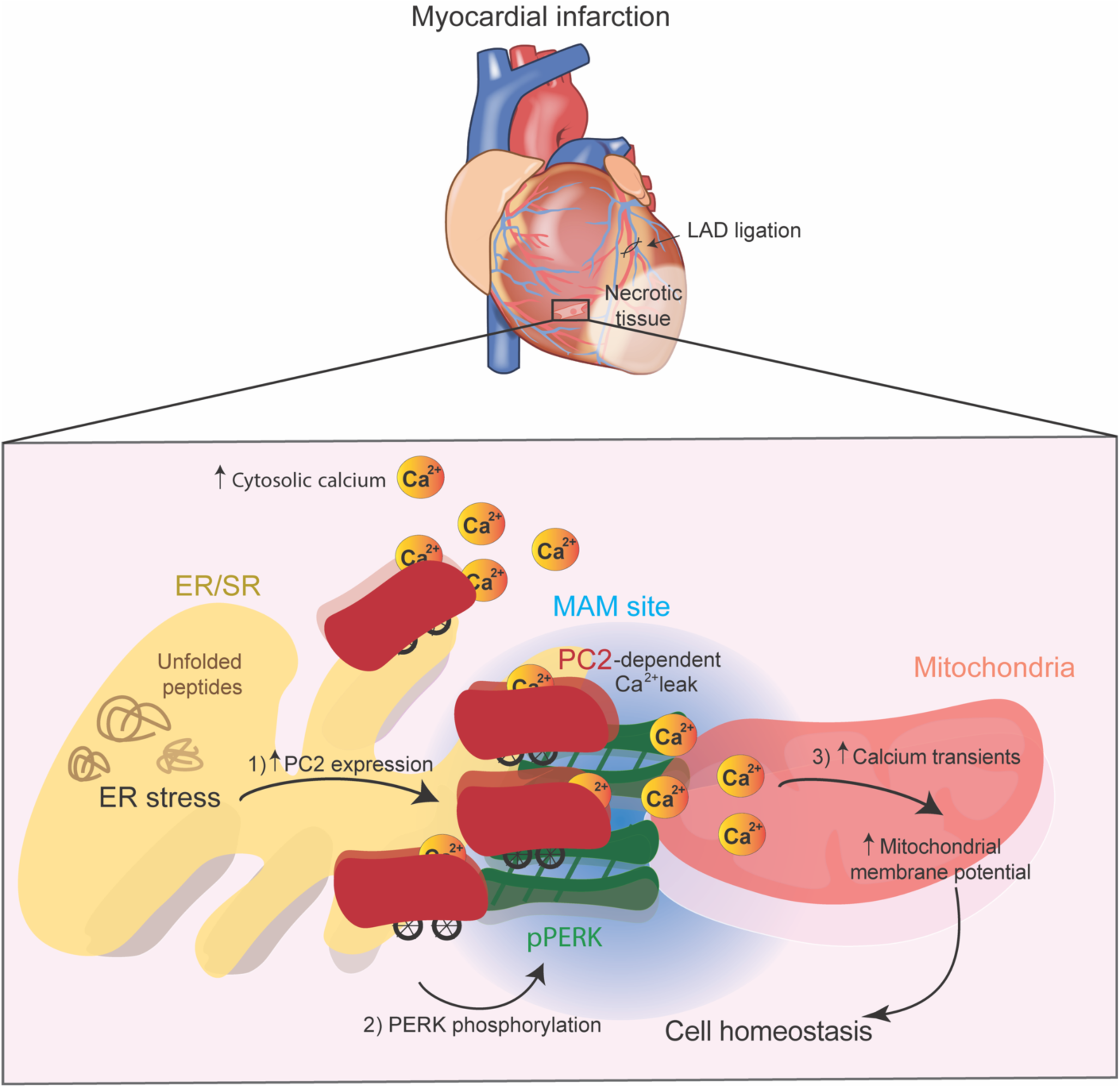

## Introduction

Cardiovascular complications are the leading cause of death in patients with autosomal dominant polycystic kidney disease (ADPKD), the most common form of genetic kidney disease, characterized by bilateral renal cyst formation^1,2^. Patients with ADPKD have an increased risk of acute myocardial infarction (AMI), worsened clinical outcome, and increased mortality post-AMI, compared to non-ADPKD patients^3,4^. However, the underlying mechanism of cardiac-related pathologies remains largely unknown. ADPKD occurs due to mutations in the *PKD1* and *PKD2* genes, which account for 80% and 12-15% of cases, respectively^1,5^. Despite being responsible for fewer cases, patients with *PKD2* mutations have an increased risk of cardiac hospitalizations, compared to patients with *PKD1* mutations^6^ and a greater prevalence of dilated cardiomyopathy (DCM) ^7,8^, which can result from AMI^7,8^. Strikingly, the largest epidemiological study of AMI in PKD patients originates from Taiwan, where ADPKD patients had a 3-fold higher prevalence of AMI and worsened outcomes^3,4^. In contrast to other populations, the Taiwanese population is predominantly affected by a *PKD2* founder mutation^9^, suggesting that mutations in PKD2 predispose individuals to AMI.

The gene products of *PKD1* and *PKD2* encode polycystin 1 and 2 (PC1 & PC2), respectively^10,11^. PC1 is an eleven-transmembrane domain protein proposed to have G protein-coupled receptor (GPCR) properties^12^. PC2, a partner protein to PC1, is a member of the Transient Receptor Potential Polycystin (TRPP) channel family, localizing to the endoplasmic/sarcoplasmic reticulum (ER/SR), plasma membrane (PM), and cilia^13–15^. These proteins are found in cardiomyocytes, with PC2 functioning both as a calcium-permeant channel and as a regulator of the ryanodine receptor 2 (RyR2)^16,17^. The release of SR calcium from RyR2 opening is tightly regulated, thereby controlling excitation-contraction (EC) coupling^18^. However, the SR/ER is known to have a calcium leak, largely attributed to RyR2, and if not managed, this contributes to calcium mishandling and cardiac failure^19^. Although a TRP channel, the conditions under which PC2 calcium flux are unclear, and PC2 has been suggested as a leak channel^20^.

One condition under which calcium leak is enhanced is ER stress^21^. Indeed, under AMI, there is increased ER stress, which is counteracted by the activation of the unfolded protein response (UPR) through the canonical PERK, IRE1, and ATF6 signaling pathways^22^. These responses are essential for cell survival as the ER/SR is responsible for the folding and post-translational modification of one-third of all proteins^23^. Unresolved ER stress and impaired UPR pathways are a hallmark of many cardiovascular diseases, including AMI^22^. PERK resides on the ER, where it becomes activated through autophosphorylation^24^, and its expression increases in a non-reperfused ischemic murine model^25^. One recently uncovered role of phosphorylated PERK (pPERK) is its regulation of cardiac mitochondrial-associated membrane sites (MAMs) under ischemic injury^26^. PERK has been shown to facilitate calcium transfer at MAMs to boost metabolism and resolve ER stress^27^. Prior studies have shown that pPERK interacts with PC2 to regulate ER stress^28^, and that PC2 is found at mitochondrial contact sites in the kidney^29^. Although PC2 expression is increased under ER stress conditions^30^, the relationship between PERK and PC2, and whether PC2, specifically its calcium channel activity, is cardioprotective under AMI by regulating ER stress, is unknown.

We therefore examined the role of PC2 in regulating ER stress during AMI and its ability to function as a calcium leak channel in the ER. Consistent with PC2 participating in ER stress, we found that PC2 knockout mice had worse cardiac function after MI and reduced UPR activation. In vitro, inducing ER stress with tunicamycin led to ER calcium leak and PERK-dependent mitochondrial calcium transients, which were diminished in PC2 knockout myoblasts. As a result, this led to a decrease in mitochondrial membrane potential and increased cell death under ER stress in PC2 knockout cells. Reintroducing functional PC2, but not the PC2 D511V mutation, rescued the cytosolic calcium leak caused by ER stress. These data demonstrate that PC2-dependent calcium leak plays a key role in AMI during ER stress.

## Materials and methods

### Human heart tissue

Human heart left ventricle samples were collected from the Loyola Cardiovascular Research Institute Biorepository and the Cleveland Clinic Biorepository. The cardiac tissue from patients with ischemic cardiomyopathy was collected during heart explantation or LVAD implantation. In healthy donor patients, cardiac tissue was obtained postmortem. The non-failing healthy donor patients did not present with a history of coronary artery disease or heart failure. All tissues were flash frozen in liquid nitrogen. Supplementary table 1 includes details over the patients history, race, age and sex.

### Animal studies

The Institutional Animal Care and Use Committee (IACUC) at Loyola University Chicago approved all animal procedures performed in this study, and the experiments were conducted by pertinent regulations and guidelines. To generate cardiomyocyte-specific *Pkd2* knockout (PC2-KO) mice, *Pkd2*^flox/flox^ mice (generous gift from S. Somlo, Yale University) were crossed with tamoxifen-inducible αMHC-MerCreMer (Jackson Laboratories) C57-Bl/6 mice as previously described^16^. αMHC-MerCreMer expressing mice fed tamoxifen were used as controls. Mice were fed with tamoxifen diet at 8 weeks of age. Both male and female mice were used in this study.

### Myocardial Infarction surgery

Mice were anesthetized with isoflurane (0.8% -1.5%) and ventilated. A left-sided thoracotomy was performed between the third and fourth ribs to expose the heart. Myocardial infarction was induced by ligating the left anterior descending (LAD) coronary artery with a single suture. The mouse was then turned onto its back, the tracheal tube was removed, and the trachea and cervical incision were sutured closed.

### Echocardiography

Mice were anesthetized with isoflurane (0.8% -1.5%) and subjected to echocardiography (Vevo2100, Visual Sonics) before and after MI. B-mode (long axis for strain analysis) and M-mode (short axis) were performed. Mice were kept on a warmed platform during the procedure and had an average heart rate of 400-500 beats/minute. Cardiac parameters were calculated utilizing the Vevo2100 VisualSonics cardiography and VevoStrain packages.

### Cell lines maintenance

C2C12 myoblast cells (purchased from ATCC) and HEK 3KO (gift from Dr. David Yule) were cultured in Dulbecco’s modified Eagle’s medium (DMEM) supplemented with 10% Fetal bovine serum (FBS) and 1% antibiotics. The cells were maintained in a 37 °C incubator with 95% air and 5% CO_2_ in a humidified environment. Previously characterized CRISPR/Cas9 Pkd2 KO C2C12 cells and non-template control cells were used^31^.

### Transfection experiments

Cells were transfected 24 hours after plating with 1-2 μg of DNA, 25 μL of PEI (concentration 1 μg/mL stock) in OPTI-MEM, and incubated for at least 24 hours. The following plasmids were used: full-length PC2 fused to mCherry (PC2mcherry), PC2 fused to mCherry with a single point mutation (D511V),^31^ gCaMP6F(cytosolic calcium indicator), R-cepia (ER calcium indicator, gift of Dr. Aleksey Zima), mito-gCaMP6F (mitochondrial calcium indicator, gift of Dr. Stefani) and split fluorescent protein-based contact site sensor (SPLICS)^32^.

### Protein extraction

For C2C12 cells, total protein was extracted using a radioimmunoprecipitation assay (RIPA) buffer (EDTA, EGTA, Triton-X, sodium deoxycholate, SDS, and NaCl, pH 8) supplemented with a protease inhibitor cocktail (Sigma-Aldrich) and phosphatase inhibitors (NaF and sodium orthovanadate, Alfa Aesar). For mouse or human protein lysates, flash-frozen tissues (left ventricles) were homogenized in RIPA buffer containing protease and phosphatase inhibitors.

### Western blotting

Protein concentration was measured using Pierce bicinchoninic acid (BCA) protein assay (Thermo Fisher Scientific). Equal amounts of protein (20-30 µg) were loaded to an SDS-PAGE gel (Bio-Rad, 4%–20% gradient) and wet transferred on a polyvinylidene difluoride (PVDF) membrane. The following primary antibodies were used: PC2 (D-3, 1:500, SC-28331; Santa Cruz Biotechnology), MFN2 (D2D10, 1:1000; Rabbit mAb; #9482S, Cell Signaling Technology), MCU (D2Z3B, 1:1000; Rabbit mAb; #14997; Cell Signaling Technology), PERK (C33E10, 1:1000; Rabbit mAb; #3192; Cell Signaling Technology), CHOP (D46F1; 1:1000; Rabbit mAb; #5554; Cell Signaling Technology), phospho-PERK (Thr980; 16F8; 1:1000; Rabbit mAb; # 3179), GRP78/BIP (11587-1-AP; 1:1000, Rabbit pAb, Proteintech Group), ATF4 (60035-1-Ig, 1:1000, Mouse mAb, Proteintech Group) and GAPDH (60004-1-Ig, 1:1000, Mouse mAb, Proteintech Group). Horseradish peroxidase (HRP)-conjugated secondary antibodies were used (Immun-Star goat anti-mouse, 1:10,000, 1705046 and Immun-Star goat anti-rabbit, 1:10,000, 1705046, Bio-rad) and activated using Clarity Max Western ECL (Bio-Rad). ChemiDoc MP imager (Bio-Rad) was used to image blots, and the protein intensity was measured using ImageLab software (Bio-Rad).

### Immunofluorescence

Murine left ventricles were excised to include ischemic, border, and remote zones and were fixed in 2% PFA for 1-2 hours and processed as previously described^16^. Murine and human tissue were embedded in OCT and 15 μm cryosections cut (LEICA CM3050 S). Then, cardiac human/murine sections or C2C12 cells were fixed with 2% paraformaldehyde (PFA) then incubated with the following antibodies PC2 (YCE2, 1:100, Mouse mAb, sc-47734; Santa Cruz Technology) & CHOP (GADD153, 1:100, Rabbit pAb, 15204-1-AP, Proteintech Group), aSMA and PERK. Secondary antibodies were Alexa Fluor 488 donkey anti-mouse IgG (1:1000, A21202, Invitrogen) and Alexa Fluor 546 donkey anti-rabbit IgG (1:800, A10040, Invitrogen). Slides were mounted with Prolong-Diamond media containing DAPI (Invitrogen) and imaged using a Zeiss 880 laser-scanning microscope with Airyscan (Zeiss, Germany). Images were processed using Zen Black software (Zeiss, Germany) and analyzed in FIJI (Version 2.16.0/1.54p, NIH). Colocalization of proteins was measured by using Mander’s 2 coefficient (Coloc2 plugin) in FIJI.

### Isolation of primary murine cardiomyocytes

Mice were anesthetized with isoflurane, and the hearts were excised, and cardiomyocytes digested as previously described^16^. Digested cells were resuspended in Tyrode’s solution containing 1 mM calcium. Cells were incubated with fluo-4 AM (2 μM, Invitrogen) and rhod-2 AM (2 μM, Invitrogen). To ensure mitochondrial loading of Rhod-2, sodium borohydride was added^33^. Cardiomyocytes were placed on a coverglass pre-coated with laminin and allowed to settle. After settling, paced cells (0.5Hz) were imaged with a sCMOS camera (Orca Flash, Hamamatsu) at 5 fps. Images were analyzed on FIJI.

### Mitochondrial membrane potential measurements

Cells were loaded with the membrane potential dye TMRE (25 nM, 20 minutes), washed, and imaged on an 880 Zeiss laser-scanning microscope with Airyscan (Zeiss, Germany). Analysis was performed using FIJI.

### Calcium imaging

Cells were transfected with gCaMP6F, R Cepia, or mGcamp6f for 24 hours. Cells were imaged in DMEM media (without phenyl red), or solutions with 0 calcium. Live-cell imaging was performed using a Zeiss fluorescence microscope. For analysis, regions were drawn on individual cells to quantify calcium in at least N=3 biological samples.

### Chemical Incubation

Cells were treated with 0.1, 0.5, 1 and 2.5 µg/ml of tunicamycin (CAS #: 11089-65-9, Cayman Chemical) for 2 hours or 8 hours for protein extraction. For calcium experiments, 24 hours after transfections, cells were live-imaged and 2.5 µg/ml of TM was added. 2 µM Thapsigargin (CAS #: 67526-95-8, Cayman Chemical) was added in transfected cells for calcium live-imaging. Cells were treated with PERK inhibitor (GSK26046414, CAS #: 1337531-36-8, Cayman Chemical) for 30 minutes, washed, and live imaged.

### Statistics

Graphs of all data were generated using GraphPad PRISM 9. As per central limit theorem, we used non-parametric distribution for n<30. Mann-Whitney U (non-parametric) was used for analyzing two independent groups, and two-way ANOVA was used for two independent groups with two factors. All graphs are shown as mean ± SEM with p-values listed in the figures.

## Results

### Cardiomyocyte localized PC2 is upregulated in human ICM patients and in a murine MI model

Mutations to PC2 are associated with a higher risk and poor outcome to AMI^4^, however, the underlying mechanisms are largely unknown. Expression of PC2, as determined by both western blot and immunofluorescent assay (IF), was upregulated in human ischemic cardiomyopathy (ICM) left ventricle (LV) samples without an ADPKD diagnosis compared to healthy donors (Figure 1A-B). Specifically, the increased PC2 expression was primarily seen in cardiomyocytes (Figure 1B). Similarly, in an MI mice model PC2 was increased both in western blot and IF (Figure 1C-D). Although all images were acquired using consistent parameters, a marked increase in PC2 expression is observed in MI samples. To visualize baseline expression levels, the images in Figure 1B and 1D have been adjusted to different brightness scales, whereas the images in Figure S1A are displayed using a uniform brightness scale for fair comparison. This increased expression was localized to the area of risk (border) zone, compared to healthy (remote) zone (FigureS1C).

**FIGURE 1:**
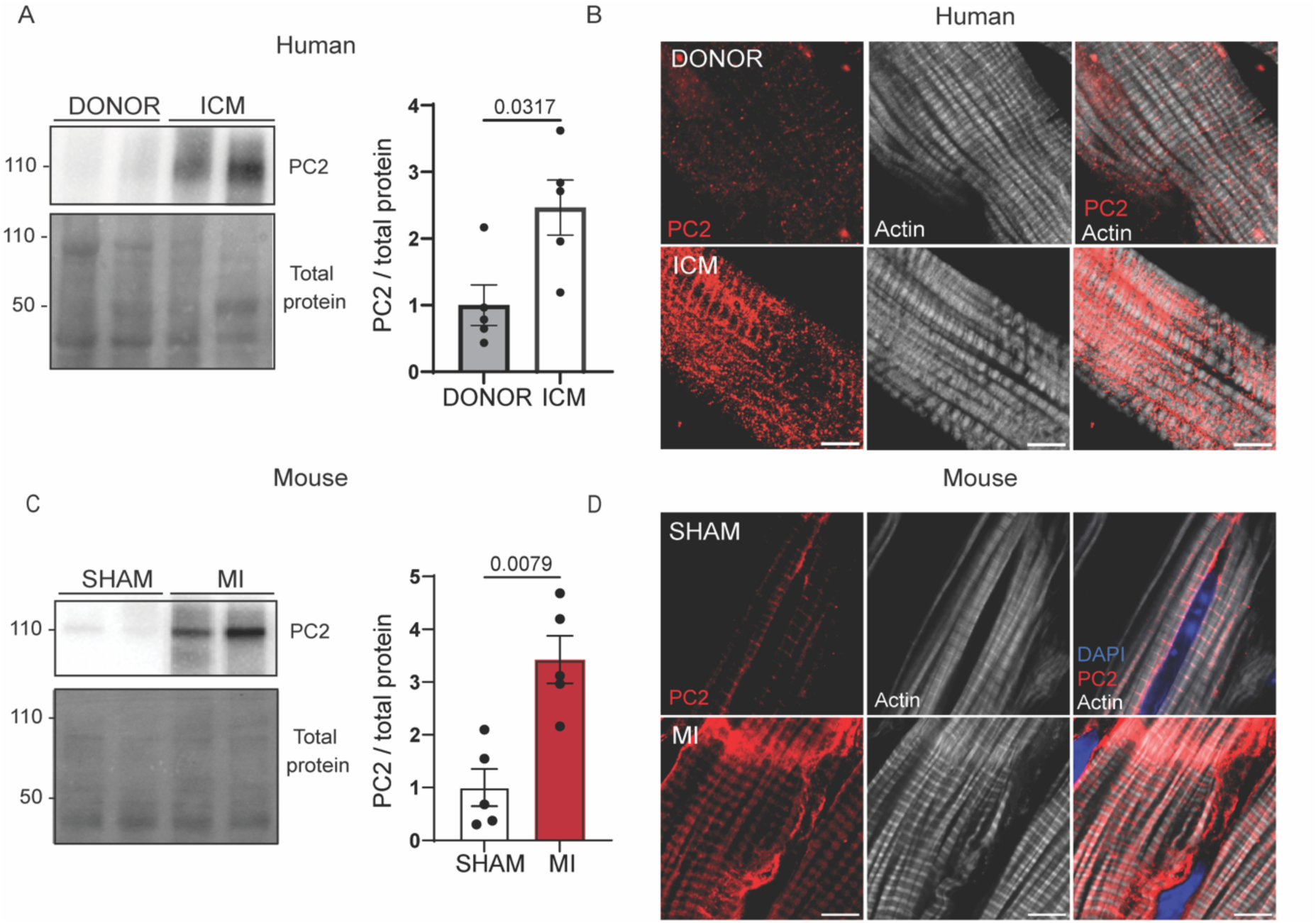
Cardiac PC2 increases in human and murine models after ischemic heart injury. **A.** Protein expression of PC2, left ventricle, from patients with ischemic cardiomyopathy (ICM) and healthy donors (DONOR), left. Total protein was used as a loading control. Quantification of western blot, right. Statistical analysis was performed using Mann-Whitney test, N=5 patients per group. **B.** Representative immunofluorescence (IF) images of PC2 (red) and actin (white) in DONOR and ICM left ventricular sections, not set to same parameters. Scale bar=5 µm. **C.** Protein expression of PC2, left ventricle, 7 days post myocardial infarction (MI) or sham (SHAM) surgery. N= 5 mice per group, left. Total protein used as loading control. Quantification of Western blot, right. Statistical analysis was performed using Mann-Whitney test. **D.** Representative IF images of PC2 (red), actin (white), and nuclei (blue) in SHAM and MI in murine left ventricular sections, not set to same parameters. Scale bar = 5µm.

### Cardiomyocyte-specific PC2-KO leads to worsened cardiac dysfunction after AMI

As PC2 was upregulated in AMI, we therefore subjected CTL and PC2-KO mice to AMI. We used a well-described cardiomyocyte-specific Pkd2 floxed model (PC2 KO), and an example IF is shown (Figure 2A). Prior studies have established that PC2 KO mice have normal systolic function at baseline^16,34,35^. In CTL and PC2-KO mice (4 weeks after deletion), MI was introduced by permanently ligating the left anterior descending (LAD) artery (Figure 2A-B). Both male and female mice were examined. As differences in MI outcomes were observed, the results are reported by sex. Using M-mode echocardiography (Figure S2A), we found that in male mice, ejection fraction (EF) was significantly decreased while systolic volume increased, post-MI in both CTL and PC2-KO mice (Figure S2B). There was no difference between genotypes. (FigureS2B). To obtain a better assessment of the left ventricular function relative to the infarct which was localized primarily to the apex (Figure 2B), we used longitudinal measurements with B-mode echocardiography to calculate cardiac strain). Overall, global longitudinal strain (GLS) significantly decreased ∼10% in CTL mice post-MI, but an even greater decrease (∼60%) was observed in the PC2-KO post-MI (Figure 2C). Time to peak analysis revealed that at the apex, the decrease in longitudinal strain was greater post-MI in the PC2-KO compared to CTL male (Figure 2D). At the mid-section, posterior longitudinal strain only decreased in the PC2-KO post-MI but not the CTL mice (Figure 2D). Longitudinal strain was unchanged at the posterior base (equivalent to a remote area post-MI, Figure 2D).

**FIGURE 2:**
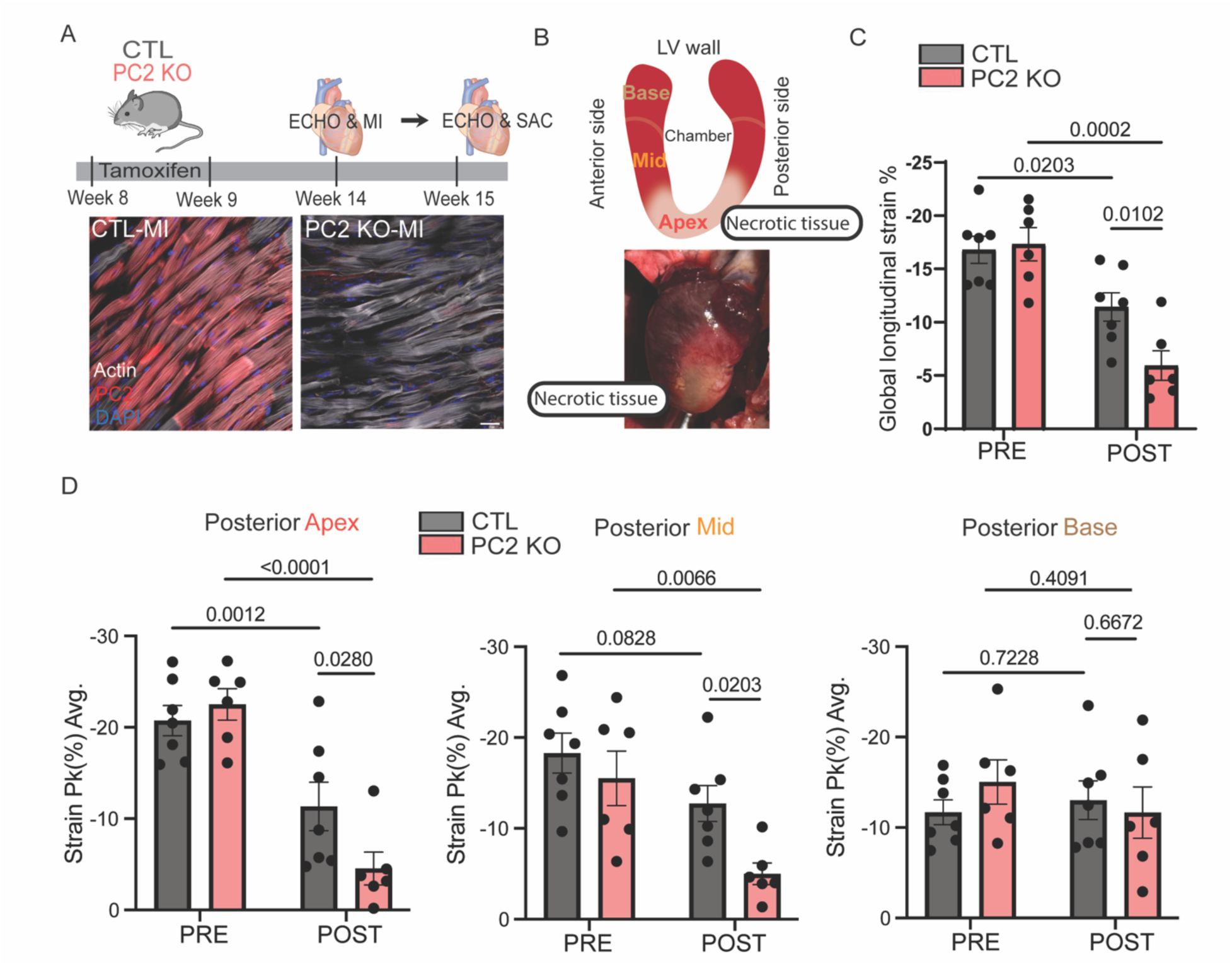
Cardiomyocyte-specific PC2 KO exacerbates cardiac dysfunction after AMI. **A.** Diagram illustrating the experimental design of the AMI surgery in the CTL and cardiomyocyte-specific PC2 KO mice. Representative IF images of PC2 (red) and actin (white) in left ventricular sections from CTL and PC2 KO mice 7 days post-MI. Images are set to the same parameters. Scale bar = 20 µm. **B.** Diagram illustrating the organization of the ventricular wall (base, mid, and apex), and the infarct area localized at the apex. Representative image of the infarct area. **C.** Global longitudinal strain (GLS) analysis in CTL (n=7, male) and PC2 KO (n=6, male) mice pre and post AMI injury. **D.** Longitudinal strain analysis of the posterior cardiac wall at the base, mid, and apex sections in CTL and PC2 KO mice pre- and post-AMI injury. Two-way ANOVA with repeated measurements was used for statistical analysis.

In contrast to the male mice, a more severe phenotype was observed in the PC2-KO female mice. EF was significantly decreased post-MI in female mice compared to their sham genetic controls. There was a further decrease in the MI PC2-KO mice compared to the MI control mice (FigureS2C). Increased heart weight was only seen in CTL female mice (post-MI) but not in PC2-KO female mice, suggestive of impaired cardiac remodeling (FigureS2C). There was a higher mortality rate in female PC2-KO mice compared to CTL mice (FigureS2D).

Overall, these data suggest that PC2 KO contributes to exacerbated dysfunction post MI. In addition, we report a sex difference in an MI model after loss of PC2 with the PC2-KO female mice being less resistant, although the cause of this increased sensitivity in female PC2 KO mice is unclear.

### Cardiomyocyte-specific PC2-KO impaired PERK-CHOP signaling and increased fibrosis

To understand mechanistically why there is worsened cardiac dysfunction in the PC2-KO mice post-MI, we examined mitochondrial function and the ER stress pathway, two well-established pathways known to be affected by and contribute to the overall outcomes of an MI^36,37^. We chose to focus on these two pathways as we noted that deletion of PC2 affected mitochondrial function and the PERK-CHOP signaling axis (FigureS3). Specifically, cardiomyocytes from PC2 KO mice (Figure S3A) had decreased mitochondrial calcium and more depolarized mitochondria compared to control cardiomyocytes (FigureS3B-C). Expression of the mitochondrial calcium uniporter (MCU) was decreased in PC2-KO mice while the mitochondrial tethering protein MFN2 was unchanged (FigureS3D-E). In contrast, there was a decrease in the expression of PERK (FigureS32F-G). PERK is a known mitochondrial tethering protein ^26^, and a constituent of the UPR stress pathway^22^. CHOP, which lies downstream of PERK in the UPR stress pathway, was significantly decreased in PC2-KO mice (Figure S3D-E).

We found that PC2, PERK, and MFN2 colocalized post MI compared to SHAM mice, suggesting a role of PC2 and PERK at the MAMs. (Figure 3A). Despite the difference at baseline in MCU expression (Figure S3D-E), MCU and MFN2 protein expression were the same between CTL and PC2-KO mice post-MI (Figure 3B-C). Post-MI PERK expression remained decreased in the PC2-KO mice compared to CTL (Figure 3D-E and FigureS5A). Additionally, there was increased expression of the pro-fibrotic marker, αSMA, around the infarct zone in the PC2-KO mice post-MI compared to CTL (Figure 3F-G). Collectively, these data indicate that loss of PC2 exacerbates cardiac dysfunction in MI by dysregulating the PERK-dependent branch of the UPR pathway and this associates with increased fibrosis.

**FIGURE 3:**
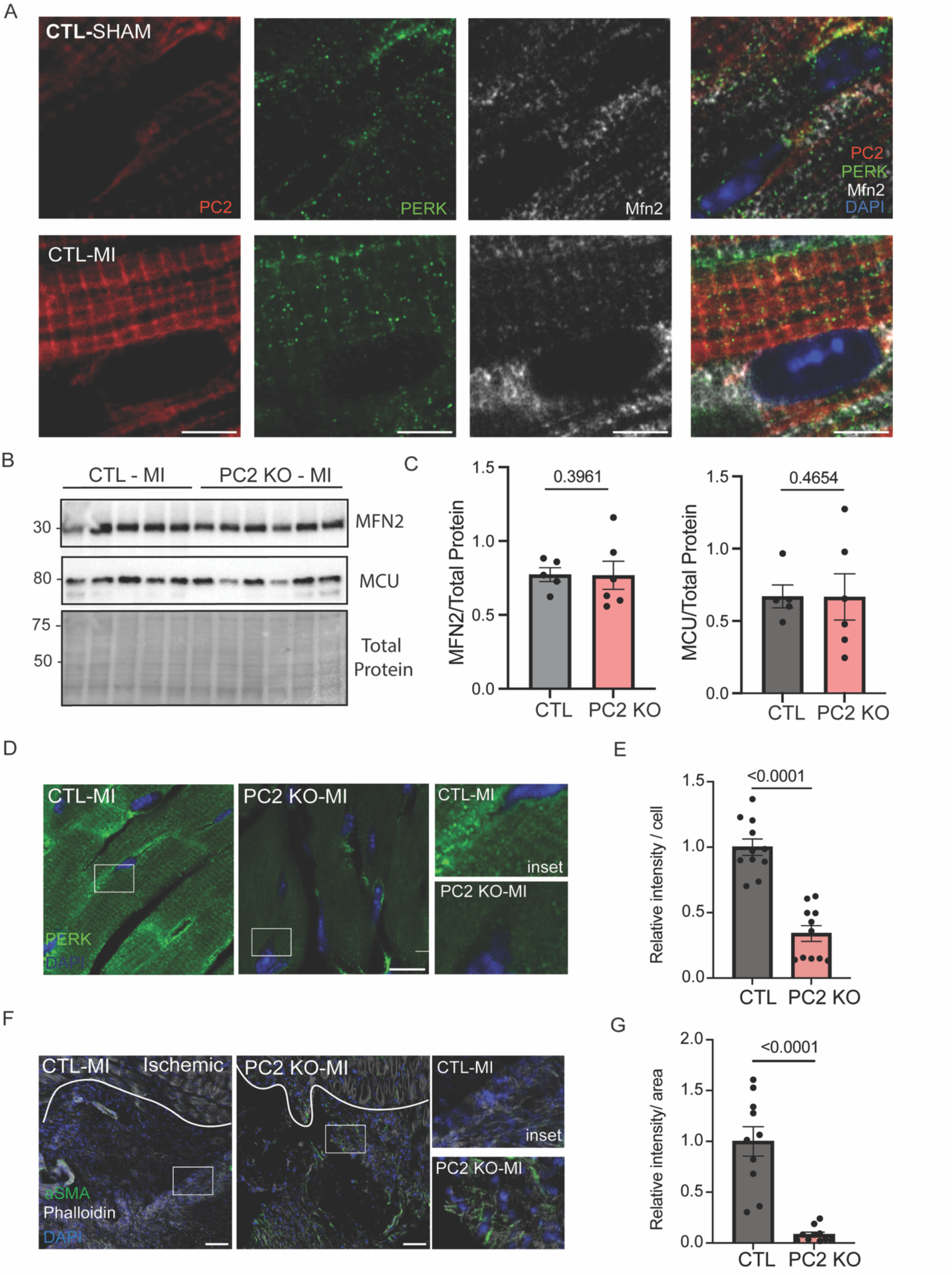
PC2-KO leads to impaired PERK-CHOP signaling and increased fibrosis. **A.** Representative IF images of PERK (green), PC2 (red), MFN2 (white) and nuclei (blue) in left ventricular sections from CTL SHAM and MI mice. Scale bar = 5 µm. **B.** Protein expression of MCU and MFN2 from CTL (n=5) and PC2 KO (n=6) mice. Total protein was used as a loading control. **C**. Quantification of A. Statistical analysis was performed using Mann-Whitney test. **D.** Representative IF images of PERK (green) and nuclei (blue) in left ventricular sections from CTL and PC2 KO male mice. Scale bar = 10 µm. Inset images taken from white square. **E.** Quantification of IF PERK expression. Each dot represents a single image with at least three cardiomyocytes from N=3 mice/group. Statistical analysis performed using Mann-Whitney test. **F.** Representative IF images from cardiac sections stained with alpha smooth actin (αSMA, green), actin (white), and nuclei (blue) in left ventricular sections from CTL and PC2 KO male mice. Scale bar 20 µm. The white line represents the division between the border and the ischemic zone. Inset images taken from the area in white square. **G.** Quantification of αSMA expression normalized to area. Statistical analysis was performed using the Mann-Whitney test. Each dot represents area from the ischemic zone in 3-4 different images from N=3 mice.

### Loss of PC2 in a myoblast cell line leads to impaired PERK-CHOP signaling axis

To better understand molecularly how PC2 is cardioprotective against MI we used a previously validated myoblast C2C12 cell line with PC2 knocked out by CRISPR/CAS9^31^. While the ER chaperone GRP78 expression remained unchanged, PERK and CHOP expressions were significantly downregulated at baseline in the PC2-KO cells (Figure 4A-B), consistent with the findings in the PC2-KO mice. To determine if these changes were seen at the transcript level, we re-examined our previously published bulk RNA sequencing dataset on PC2 KO cells^31^. We found that CHOP was significantly down-regulated in PC2-KO cells (Figure 4C). However, PERK gene expression remained unchanged, indicating that PC2 regulates PERK at the translational level.

**Figure 4:**
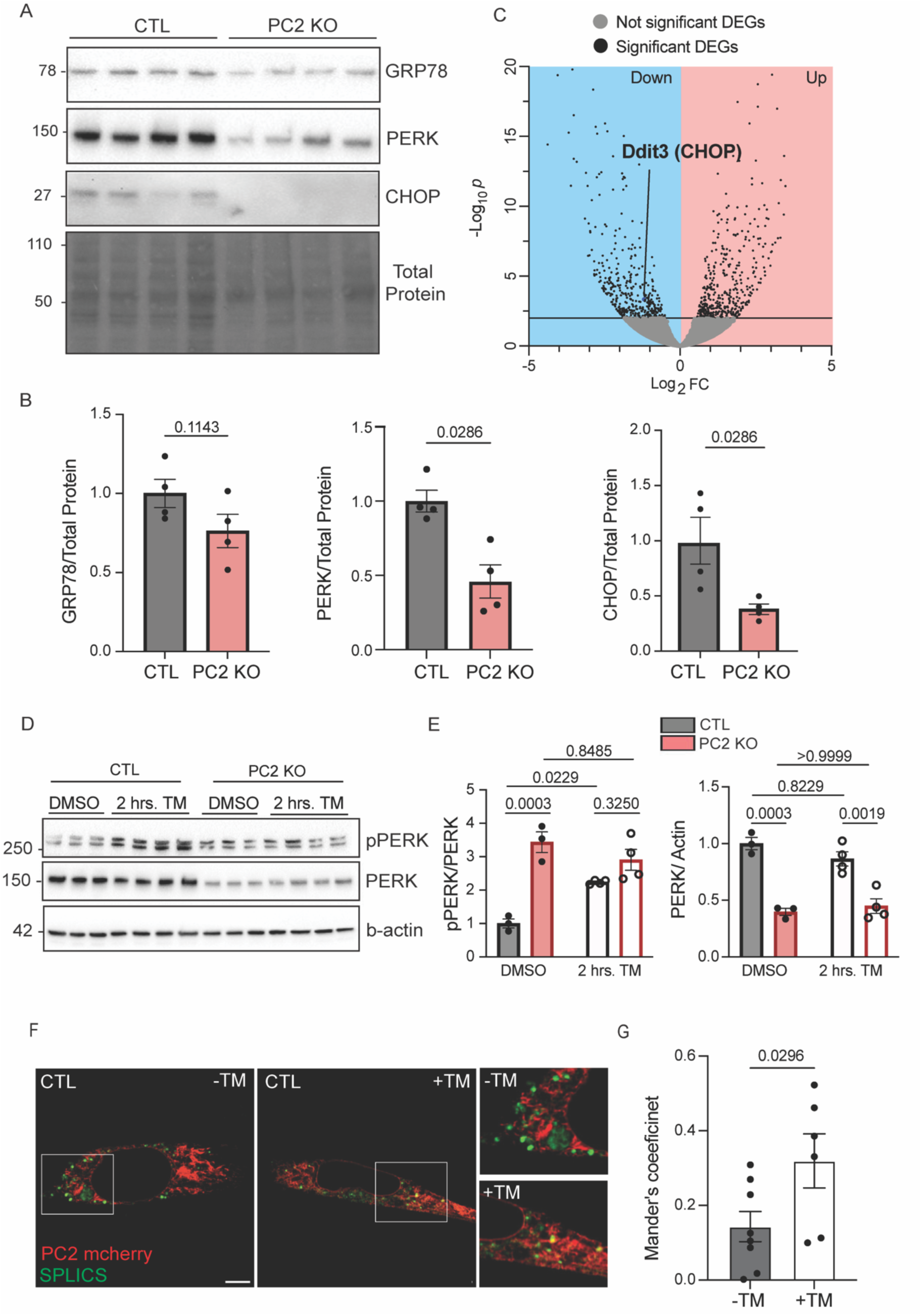
PC2 regulates expression of PERK and CHOP and localizes close to MAM sites under ER stress. **A.** Protein expression of GRP78, PERK and CHOP in CTL and PC2 KO C2C12 cells. Total protein used as loading control. N=4 biological replicates. **B.** Quantification of A. Statistical analysis was performed using Mann-Whitney test. **C.** Volcano plot of differentially expressed genes (DEGs) upregulated and downregulated in PC2 KO cells, compared to CTL cells. N=3 biological replicates. **D**. Protein expression of PERK from CTL and PC2 KO cells treated with DMSO (vehicle) or 2.5 µg/ml tunicamycin (TM) for 2 hours. GAPDH used as a loading control. **E.** Quantification of D. N=3-5 biological replicates. Statistical analysis was performed by using a two-way ANOVA test. **F.** PC2-mcherry (red) and SPLICS (green) were transfected in CTL cells, and representative images of cells were taken before (left panel) or 30 minutes (mid panel) after the addition of TM, 2.5 µg/ml. Representative insets (right panel) images taken from the area in white squares. **G.** Mander’s coefficient analysis of the colocalization of ER-mitochondrial membrane sites (ER splics, green) and PC2-mcherry (red).

As PERK can function as an ER stress sensor, and MI causes ER stress, we tested the effect of chemical ER stress. C2C12 cells were subjected to 2.5 µg/mL of tunicamycin for 2 hours. Tunicamycin (TM) blocks N-linked glycosylation and leads to the accumulation of unfolded proteins in the ER^38^. Consistent with an activation of ER stress, p-PERK was increased after TM treatment in CTL cells (Figure 4D-E). Although p-PERK was higher in PC2 KO cells compared to the CTL cells at baseline, there was no effect of TM (Figure 4D-E). Total PERK expression remained decreased in the PC2-KO cells compared to CTL cells after TM treatment (Figure 4D-E). To test if the PC2-KO cells were able to sense ER stress, we conducted a dose-dependent titration of TM treatment for 8hrs in both CTL and PC2-KO cells. We examined the expression of GRP78, which is upstream of PERK, and the downstream targets of the PERK signaling pathway, CHOP and ATF4. In the CTL cells, there was a dose-dependent increase of GRP78, ATF4, and CHOP with increased concentrations of TM (0.1, 0.2 0.5, 1 and 2.5µg/mL) (Supp. Fig. 4A & C). However, in the PC2-KO cells, there was repressed up-regulation of GRP78, ATF4 and CHOP in response to increasing doses of TM treatment (FigureS3B-D). These data suggest that PC2 KO cells have a decreased level of PERK at baseline and a blunted response to ER stress.

As PERK is also a tethering protein, localizing at MAMs^26^, and PC2 can interact with PERK^28^, we tested whether PC2 resides at the MAMs under ER stress. To assess this, we expressed PC2-mCherry in CTL cells together with a split-GFP-based ER-mitochondrial contact site sensor (SPLICS), which fluoresces when the ER and mitochondria form MAM sites^32^ (8-10nm in proximity) (Figure 4F). After 30 minutes of TM treatment, we found an increased colocalization of PC2-mCherry with the SPLICS sensor (Figure 4F). The increased co-localization between PC2 and SPLICS indicates that PC2 moves to the ER-mitochondrial contact sites under ER stress. Altogether, these data show that the level of PC2 in the cell regulates expression and phosphorylation of PERK, and upon ER stress, PC2 becomes enriched at MAM sites.

### ER localized PC2 acts as a leak channel, mediating mitochondrial calcium under ER stress

The MAM sites enable the effective transfer of calcium from the ER to the mitochondria, facilitating ATP production, which is increased under ER stress conditions, thereby sustaining the folding capacity in the ER^21,39^. As PC2 moved to MAM sites with ER stress (Figure 4F), and PC2 is a calcium permeant channel, cytosolic and mitochondrial calcium were measured with gCaMP6F (cytosolic calcium), R-cepia (ER-calcium), or mito-gCaMP (mitochondrial calcium). In experiments where ER calcium and cytosolic calcium were measured simultaneously, TM induced an ER calcium release (decrease of R-cepia fluorescence) that coincided with a slow increase in cytosolic calcium in CTL cells (Figure 5A). The slope of the rising phase of cytosolic calcium was higher in CTL than in PC2 KO cells (FigureS5B). These experiments were conducted with 0 mM extracellular calcium to ensure that the cytosolic calcium leak originates from intracellular calcium release. However, in PC2 KO cells, TM treatment induced a blunted ER calcium release and unchanged cytosolic calcium in PC2-KO cells (Figure 5B-C). To test that the decreased calcium in response to TM was not due to decreased ER calcium store in PC2 KO cells, we used thapsigargin (Tg) and found no difference in the amount of ER-calcium storage between CTL and PC2 KO cells (FigureS5E). These data suggest that PC2 is involved in the ER calcium leak upon TM treatment. Several molecules have been associated with the ER leak^20^. We tested whether the IP_3_Rs contributed to the leak by examining the effect of TM in HEK cells with the three isoforms of the IP_3_Rs knocked out^40^. In the 3KO-HEK cells, we observed calcium leak after TM treatment (FigureS5B-C). Another potential source of the leak is the translocon, which is regulated by GRP78 (also known as BIP)^41^. However, we found that GRP78 levels were unchanged in PC2 KO cells following ER stress (FigureS5D), suggesting that GRP78 is not responsible for the blunted calcium leak we see in PC2 KO cells treated with TM. These data show that PC2 is a likely source of the calcium leak following ER stress.

**Figure 5:**
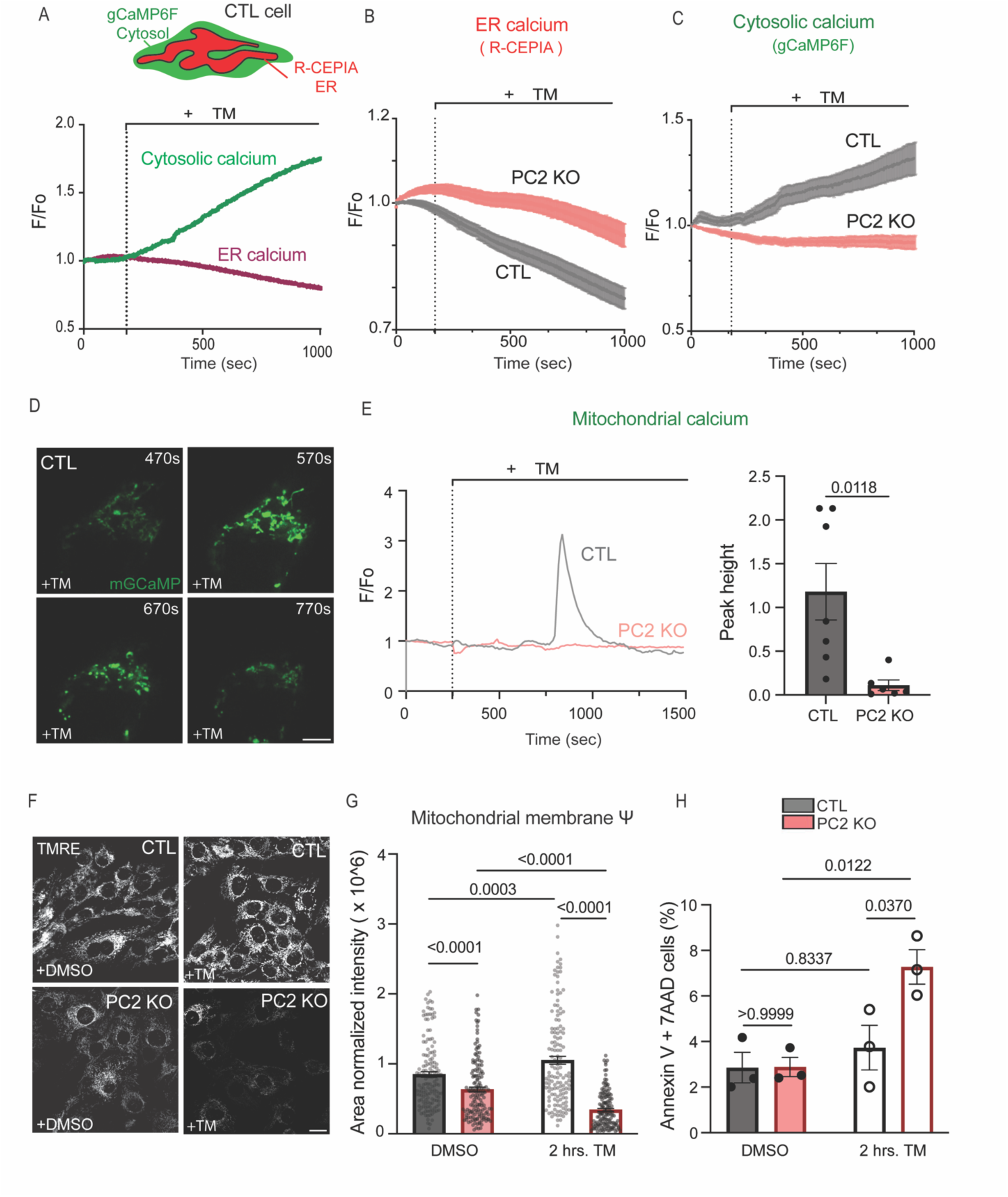
PC2 promotes calcium release from the ER and mitochondrial transients under ER stress to protect against cell death. **A.** Diagram of different genetic calcium indicators used for the ER and the cytosol. Representative traces of ER (R-cepia, red) and cytosolic calcium (GCaMP6f, green) in CTL cells treated with TM, 2.5 µg/mL **B.** Averaged ER calcium (R-cepia) traces in CTL and PC2 KO cells treated with TM, 2.5 µg/mL **C.** Averaged cytosolic calcium (GCaMP6f) traces in CTL and PC2 KO cells treated with TM, 2.5 µg/mL. N=20-23 cells from CTL and PC2 KO cells from 3 independent experiments. N=20-27 cells from CTL and PC2 KO cells from 3 independent experiments. **D.** Time course representative image of mitochondrial GCaMP (mitoGCaMP) in CTL cells treated with TM, 2.5 µg/mL. Scale bar = 5 µm. **E.** Representative traces from CTL and PC2 KO cells expressing mitoGCaMP and treated with TM, 2.5 µg/mL, left. Peak height analysis of mitochondrial calcium transients, right. Statistical analysis was performed by the Mann-Whitney test from 3-5 independent experiments. **F.** Representative images of CTL and PC2 KO cells loaded with TMRE to measure mitochondrial membrane potential (λλ′) after 2 hrs. of treatment with DMSO (vehicle) or TM, 2.5 µg/mL. Scale bar =20 µm. **G.** Analysis of mitochondrial membrane potential in CTL and PC2 KO cells treated for 2 hrs. with DMSO or TM, 2.5 µg/mL. Statistical analysis was performed by using a Two-way ANOVA from 3-4 independent experiments. **H.** Analysis of annexin-5 and 7AAD staining following treatment for 2 hrs with DMSO or TM, 2.5 µg/mL. N=3 biological replicates. Statistical analysis was performed by using two-way ANOVA from 3 independent experiments.

As we found that PC2 becomes enriched at MAM sites following ER stress, we investigated whether calcium is transferred into the mitochondria. TM treatment led to a transient rise in mitochondrial calcium in CTL cells, which was not seen in PC2-KO cells (Figure 5D-E). Notably, unlike other reported increases in mitochondrial calcium with tunicamycin^27^, the rise was of a transient nature and not sustained (not indicative of mitochondrial calcium overload). As transient rises in mitochondrial calcium can increase mitochondrial metabolism, we tested mitochondrial membrane potential 2 hours after TM treatment. Consistent with the mitochondrial calcium transients observed in the CTL cells, mitochondrial membrane potential increased after TM treatment (Figure 5F-G). Compared to CTL cells, the mitochondrial membrane potential in PC2-KO cells was more depolarized at baseline and became even more depolarized after TM treatment (Figure 5G). Consequently, PC2-KO cells exhibited increased apoptosis after ER stress induction. In contrast, the CTL cells did not have increased apoptosis after TM treatment (Figure 5H).

### PC2 channel activity is necessary for cytosolic calcium increase under ER stress

To test if PC2 channel activity was necessary for the TM induced calcium leak, we re-expressed wild-type PC2-mCherry or the D511V mutant. PC2-D511V is a pathological mutation in the voltage-sensing region that impairs ion channel activity, while retaining ER localization^42^. Re-expression of PC2-mCherry in PC2-KO cells resulted in a slow rise of cytosolic calcium, consistent with the calcium rise in CTL cells after TM treatment, and a comparable slope to CTL cells. (Figure 6A, *red line*). Importantly, re-expression of the D511V mutant version did not rescue the calcium signal (Figure 6A, green line), confirming that PC2 channel activity is required for the TM-induced ER calcium leak.

**Figure 6:**
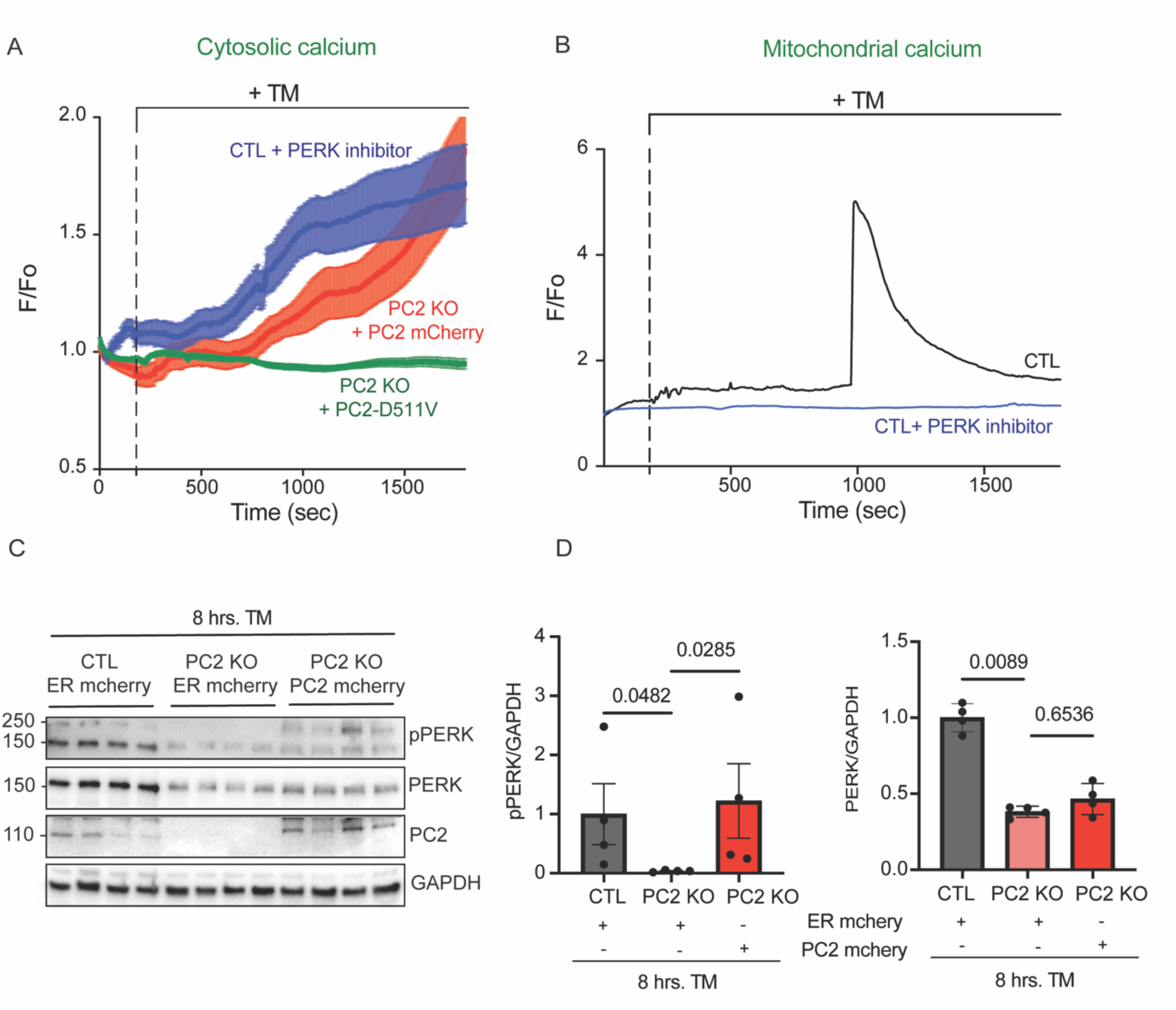
PC2 channel activity is necessary for the cytosolic calcium increase after ER stress. **A.** Averaged traces of cytosolic calcium (GCaMP6f) from CTL cells pre-treated with 1 µM PERK inhibitor for 30 minutes (blue trace) and PC2 KO cells expressing either PC2-mCherry (red trace) or PC2-D511V mutant (green trace) treated with TM, 2.5 µg/mL. **B.** Representative traces of mitochondrial calcium from CTL cells (black trace) or CTL cells pre-treated with 1 µM PERK inhibitor for 30 minutes (blue trace) then treated with TM, 2.5 µg/mL. **C.** Protein expression of pPERK, PERK, PC2, CHOP in CTL cells expressing ER-mCherry, PC2 KO cells expressing ER-mCherry or PC2-mCherry. GAPDH was used as a loading control. N=4. **D.** Quantification of pPERK and PERK from C. Statistical analysis was assessed by using one-way ANOVA.

Taken together, these data support the notion that PC2 functions as an ER-stress-activated calcium leak channel during the adaptive phase of ER stress, promoting mitochondrial calcium uptake to enhance metabolism. Conversely, loss of PC2 leads to increased cell death under ER stress.

### PERK phosphorylation is required for the transfer of calcium to the mitochondria even in the presence of PC2

We had observed that the PC2 KO cells had decreased PERK levels, and a decreased ability to be phosphorylated. To determine if PERK activation (phosphorylation and dimerization) contributes to the calcium leak from the ER, we used a selective PERK inhibitor (GSK2606414) and pre-treated cells with 1 mM for 30 mins prior to TM addition. PERK inhibition did not affect the cytosolic calcium rise after treatment with TM (Figure 6A, *blue line*). However, the inhibition of PERK lead to a blunted mitochondrial calcium response (Figure 6B, blue line, and FigureS5A). These results suggest that PERK activation is needed to relay the cytosolic calcium signal to the mitochondria. We tested whether re-expression of PC2 was sufficient to restore PERK expression in the PC2-KO cells. Re-expression of full-length PC2 restored the ability of PERK to be phosphorylated but not the total PERK expression in PC2-KO cells after TM treatment (Figure 6C-D).

### ER stress induced calcium leak in CTL cardiomyocytes and increased PC2 expression

We showed in myoblast cells that TM-induced ER stress is mediated by PC2. To test if this also held true in cardiomyocytes, we examined cytosolic and mitochondrial calcium in isolated adult cardiomyocytes from CTL and PC2 KO mice. Notably, the SR calcium stores were unchanged in PC2 KO cardiomyocytes compared to CTL^16^.

To test if ER stress induction similarly causes PC2-dependent calcium rise in cardiomyocytes, we measured paced cytosolic calcium transients following acute TM application (Figure7A). While the overall amplitude of the signal was unchanged between CTL and PC2-KO after TM application, we found a rise in diastolic calcium in CTL cells, not seen in the PC2-KO cells (Figure7B). There was a rise in mitochondrial calcium in CTL cells following TM treatment, whereas PC2 KO cells had a decreased change in mitochondrial calcium (Figure 7E-D). As ER stress also causes calcium alternans, we were not surprised to note calcium alternans (Figure7C). These calcium alternans are either due to the phosphorylation of RyR or the loss of RyR inactivation by calmodulin^19^. We found that 75 % of CTL cardiomyocytes had increased propensity for calcium alternants compared to only 46% in PC2-KO cardiomyocytes (FigureS6A). Finally, we tested if acute ER stress was sufficient to drive the upregulation of PC2, as seen in human and murine ischemic hearts (Figure 1). TM treatment induced upregulation of PC2 in both cardiomyocytes (Figure 7F) and myoblast cells (FigureS6B).

**Figure 7:**
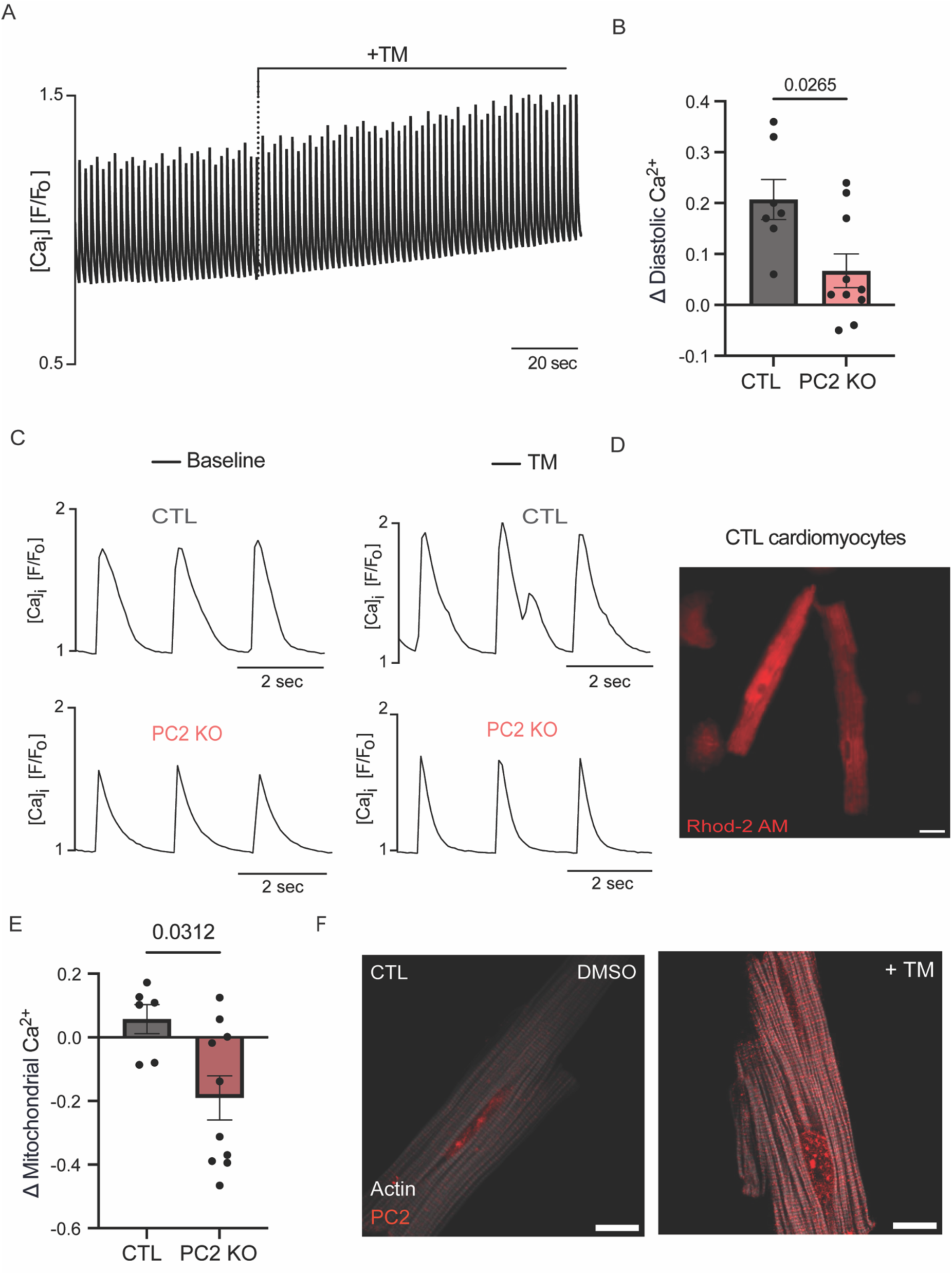
PC2 expression increases in cardiomyocytes after TM induced ER stress, and TM induces a diastolic rise in isolated cardiomyocytes. **A.** Representative cytosolic calcium (fluo4) with 0.5Hz pacing of a CTL cardiomyocyte treated with 2.5 µg/mL TM**. B.** Diastolic calcium measurement after TM treatment in CTL and PC2 KO cardiomyocytes. Each dot represents the delta change in diastolic calcium of a cardiomyocyte before and after TM treatment. Statistical analysis was assessed by using Mann-Whitney test. N=4 animals/group. **C**. Representative calcium (fluo4) transients from CTL and PC2 KO cardiomyocytes with or without TM 2.5 µg/ml. Example traces are taken from the same cell. **D.** Representative images of the mitochondrial calcium indicator Rhod2 in CTL cells. **E.** Mitochondrial calcium measurement following TM treatment in CTL and PC2 KO cardiomyocytes. Each dot represents the delta change in mitochondrial calcium in a cardiomyocyte before and after TM treatment. Statistical analysis was assessed by using Mann-Whitney. N = 4 animals/group. **F.** Representative IF images of PC2 (red) and actin (white) of CTL cardiomyocytes treated with DMSO or TM, 2.5 µg/mL, for 30 minutes.

## Discussion

The aftermath of an AMI requires cardiomyocytes to mount stress responses to counteract the infarct^22,43^. In this study, we demonstrate that polycystin 2 (PC2) has three key features that make it a cardioprotective molecule. First, PC2 protein expression increases in both human and murine ischemic hearts, and the loss of cardiac-specific PC2 during AMI leads to worsened cardiac dysfunction, along with dysregulated UPR response. Second, we show that PC2 functions as an ER calcium leak channel under ER stress. Third, we show that PC2 becomes enriched at ER-mitochondrial sites where its expression regulates in activation of PERK, aiding mitochondrial calcium transfer to enhance metabolism and restore cellular homeostasis. Overall, our data identify PC2 as a new stress sensor that is dynamically upregulated by ER stress and relocates to MAMs to integrate both the UPR and intracellular calcium signaling in the context of AMI.

### PC2 is a leaky calcium channel under ER stress

Under pathological conditions, including AMI, there is an increased calcium leak from the SR^44^. This has been variously attributed to the RyR, IP_3_Rs, and the translocon SEC61^19,41,45^. (REF, missing IP3). Enhanced RyR leakiness can arise from its phosphorylation or from interactions with calmodulin^46^ (Either way, inhibition of the RyR leak has been shown to be cardioprotective^47^. However, the leaky nature of the ER/SR can also be beneficial, as it allows sufficient calcium to be taken up by the mitochondria, thereby enhancing metabolism^27,48–50^. However, the molecules that contribute to the beneficial calcium leak in cardiomyocytes remain unknown.

Here, we show that the dynamic upregulation of PC2, a well-defined calcium permeant channel, under conditions of ER stress fulfills the criteria of providing a “leak” that is not RyR dependent. In non-RyR expressing cells, we demonstrate that PC2 by itself, independent of the InsP3R causes a leak of calcium with ER stress, which is largely attenuated when PC2 is KO. Importantly, re-expression of PC2 but not the D511V mutant (PC2 dead pore) was able to rescue the slow leak of calcium induced with ER stress. In cardiomyocytes, we show that ER stress causes a rise in diastolic calcium that is not seen in PC2 KO cardiomyocytes. We suggest that this diastolic calcium can be transferred to the mitochondria to boost metabolism. At the same time, the rise in diastolic calcium likely lowers the threshold for RyR openings, thus resulting in calcium alternans (FigureS6). Taking our data in conjunction with prior studies, this suggests that inhibiting the RyR leak is beneficial, as it lowers calcium alternans; however, the rise in diastolic calcium contributed to by PC2 calcium leak is cardioprotective.

### PC2 and PERK interplay to resolve ER stress

AMI also results with ER stress, which if unresolved can contribute to heart failure^22^. PERK is one of the three major signaling branches of the UPR pathway, which in its active form has been shown to interact with PC2 ^28,51^. Although cardiac function is unchanged at baseline in cardiac knockouts of PERK and PC2, others have shown that PERK plays an indispensable role in regulating cardiac stress^52,53^. In our study, we demonstrate that PC2 is cardioprotective against ischemia by regulating the PERK signaling axis induced by ER stress. Curiously, some studies have suggested that PERK is protective against ischemic injury and cardiac hypertrophy ^52,53^, while others suggest that loss of PERK is cardioprotective^25^. These disparate findings suggest that PERK acts as double edge-sword depending on the conditions. Indeed, PERK activation in the initial phases of ER stress inhibits global translation to decrease the folding load, while also allowing the translation of key genes that aid in restoring cellular homeostasis^23^. However, if the ER stress cannot be resolved, the PERK signaling axis switches toward a maladaptive apoptotic pathway^23^.

One place where PERK has been found is at MAMs, where it promotes calcium transfer to maintain bioenergetics under the adaptive ER stress^27^. However, since PERK is not a channel, another protein must be responsible for the calcium flux. This role has been attributed to the InsP3R in non-cardiomyocytes and the RyR in cardiomyocytes, where, with high levels of ER stress, a sustained mitochondrial calcium level was observed, consistent with mitochondrial calcium overload. In contrast to these elevated levels of ER stress, we find that a low dose of tunicamycin is sufficient to induce a transient rise in mitochondrial calcium and hyperpolarization-conditions which are akin to the adaptive phase of ER stress. Under these conditions, we demonstrate that PC2 is a key regulator of PERK expression and activation. Indeed, loss of PC2 resulted in decreased PERK levels, and re-expression of PC2 resulted in PERK phosphorylation. PERK activation is shown to be calcium dependent^54^; thus, one interpretation of this data is that under ER stress, PC2-induced calcium leak mediates PERK activity. How PERK may be sensing ER calcium levels or a leak is beyond the scope of this study. As PC2 is known to interact with proteins in protein quality control and the UPR, which is tightly coupled to ERAD (endoplasmic reticulum-associated degradation), we may postulate that the decreased expression of PERK is due to the loss of its stability in the absence of PC2. Nonetheless, our data places PC2 as a calcium channel at the MAMs that is dynamically upregulated with ER stress, and together with PERK as a scaffolding protein, provides the calcium for mitochondria to enable the early adaptive response to ER stress and restore cell homeostasis.

## Conclusion

We identify a novel cardioprotective role of PC2 against AMI by mediating PERK signaling and calcium response during ER stress. Although the loss of PC2 has been linked to cardiovascular diseases, its role in cardiomyocytes remains largely unknown. This study highlights how PC2 helps cells adapt to ER stress to prevent cardiac injury. It also shows that PC2 functions as a leaky channel in cardiomyocytes, and these findings add to the limited understanding of this mechanism, particularly regarding when calcium leak can be beneficial for cardiac adaptation to stress.

## Novelty and Significance

### What is Known?

- PC2 is a calcium channel expressed in the heart, and patients with mutations in the PC2 gene have an increased risk and worse outcomes after MI.
- MI is characterized by dysregulated calcium handling, calcium leak, mitochondrial dysfunction, and impaired UPR signaling.
- PC2 regulates mitochondrial bioenergetics in non-cardiac cells, and PC2 increase protects against cell death by regulating the UPR.
- Under the adaptive phase of ER stress, mitochondrial calcium is necessary to rescue the cell from maladaptive signals and cell death

### What new information does this article contribute?

- PC2 increases in cardiomyocytes in the ischemic hearts of humans and mice where it can function PC2 acts as a calcium leak channel under conditions of ER stress
- PC2 is enriched at the mitochondria under ER stress where together with the scaffolding and ER-stress protein PERK, it provides a source of mitochondrial calcium to boost metabolism and restore homeostasis.
- When subjected to MI, PC2 KO mice have worsened cardiac outcome and decreased PERK expression. Our study provides evidence that PC2 is the channel that regulates the interplay between the ER and mitochondria to rescue the cell from ER stress.

## Acknowledgements

We thank Dr. Sala, and members of the Kuo lab for helpful discussion. For reagents, we thank Dr. David Yule (University of Rochester, HEK 293 TKO cells), Dr. Aleksey Zima (Loyola University Chicago, R-cepia) and Dr. Stefan Somlo (Yale University, *Pkd2* floxed mice). We thank Dr. Jordan Beach for access to the imaging resource center (Zeiss 880-Airyscan).

## Source of funding

This work was supported by National Institute of Diabetes and Digestive and Kidney Diseases (U2CDK129917 and TL1DK132769), National Heart, Lung, and Blood Institute (T35HL120835) and the Office of the Director (1S10OD028449). IYK: National Institute of Diabetes and Digestive and Kidney Diseases (R00DK101585) and PKD Foundation (1021282). VV: American Heart Association (23PRE1026740).

## Authors’ disclosure

JAK provided consulting and conducted collaborative studies with various pharmaceutical companies, but all such work is unrelated to the content of this manuscript. No other disclosures reported.

## Notes

### Competing Interest Statement

J.A.K. provided consulting and conducted collaborative studies with various pharmaceutical companies, but all such work is unrelated to the content of this manuscript. No other disclosures reported.

