## supplementary data for "Polycystin-2 is cardioprotective against myocardial infarction by regulating the calcium-mediated ER stress response"

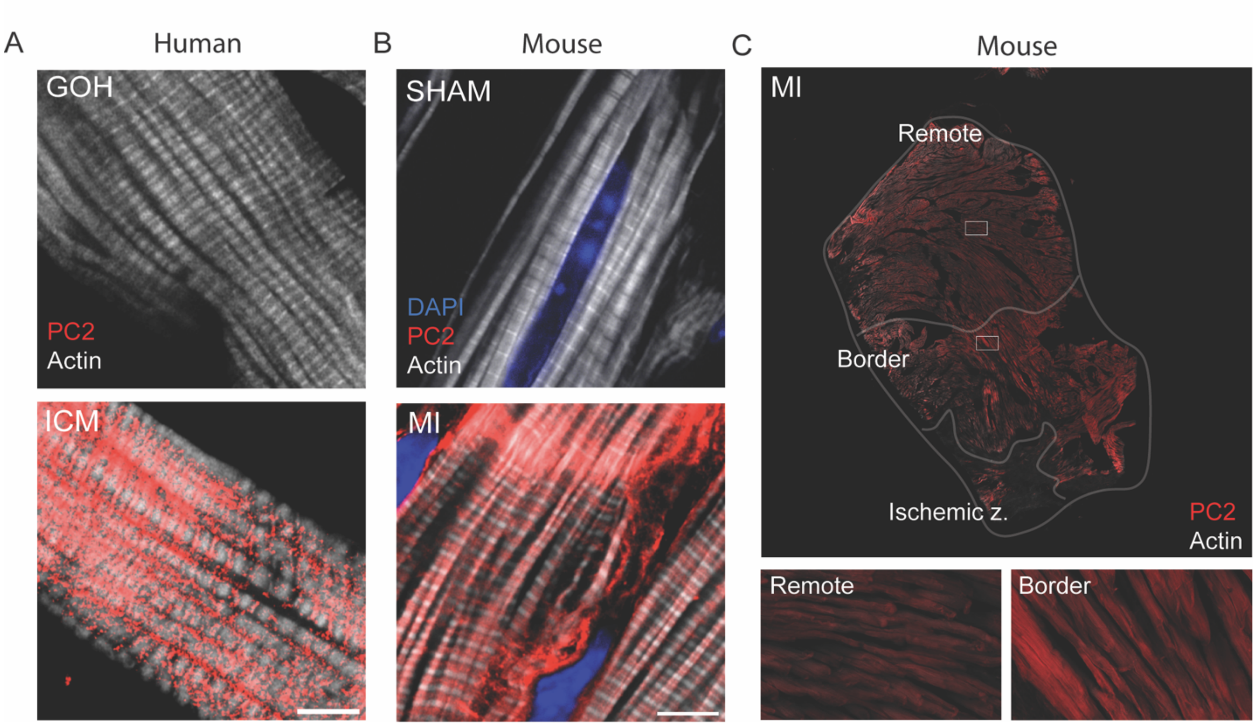


**Supplementary Figure 1. A.** Representative IF of PC2 (red) and actin (white) in DONOR and ICM human left ventricular sections. Scale bar=5 μm. **B.** Representative IF of PC2 (red) and actin (white) staining in SHAM and MI murine left ventricular sections. Scale bar=5 μm. **C.** Representative IF image of PC2 (red) and actin (white) of the left ventricle in a CTL mice 7 days post MI. Insets on right demonstrate remote (healthy) zone and border (area of risk) zone.

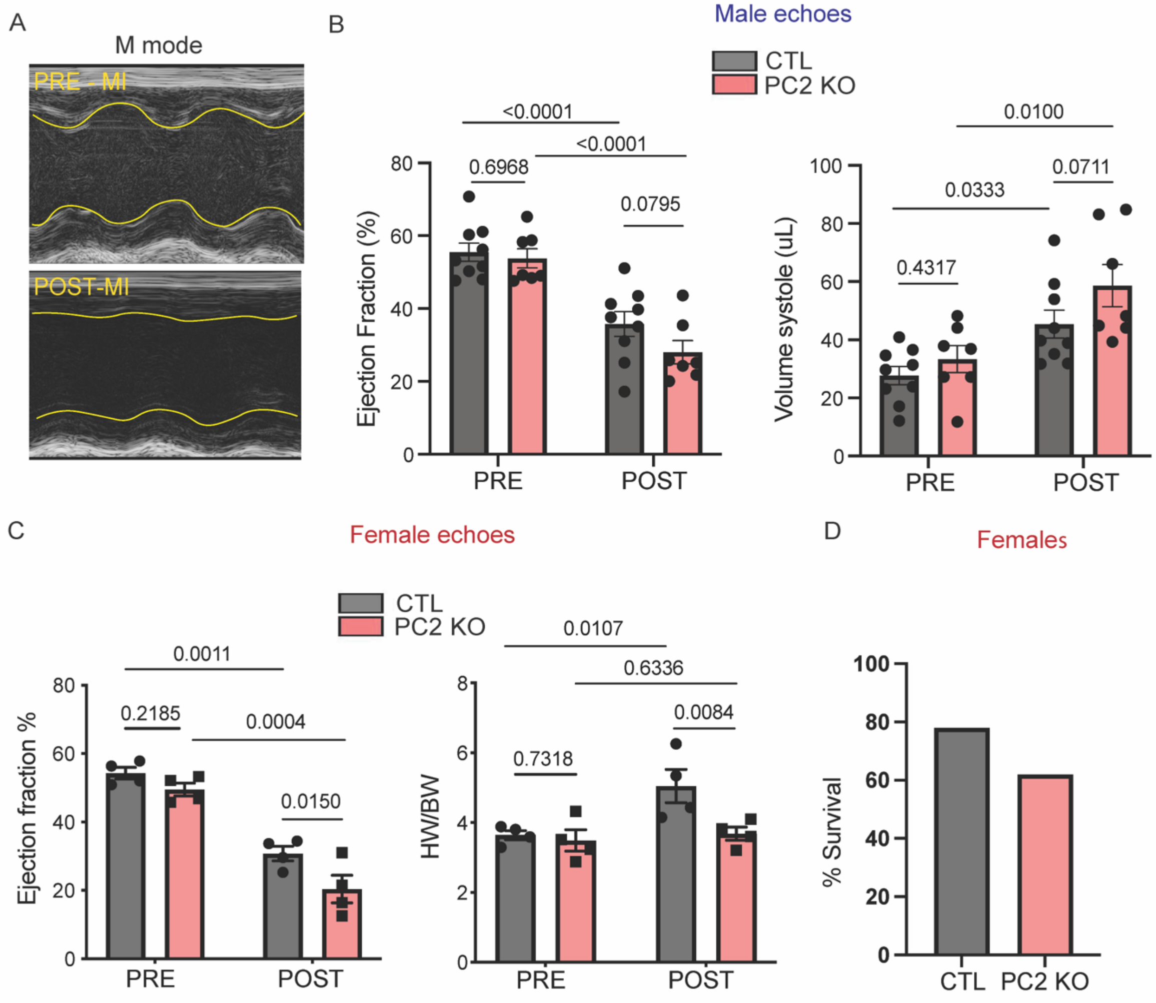


**Supplementary Figure 2: A.** Representative M-mode echo images from a CTL mouse pre-MI and 7 days post MI. **B.** Ejection fraction and systolic volume measurements of CTL and PC2 KO male mice pre and 7 days post MI (N=7-8 animals per group). **C.** Ejection fraction and heart weight measurements of CTL and PC2 KO female mice pre and 7 days post MI (N=4 animals per group). **D.** Survival (at time of MI surgery) analysis in CTL and PC2 KO female mice (N=14 animals per group). Statistical analysis in all was assessed by using Two-Way ANOVA with repeated measurements from N=4-5 animals/group.

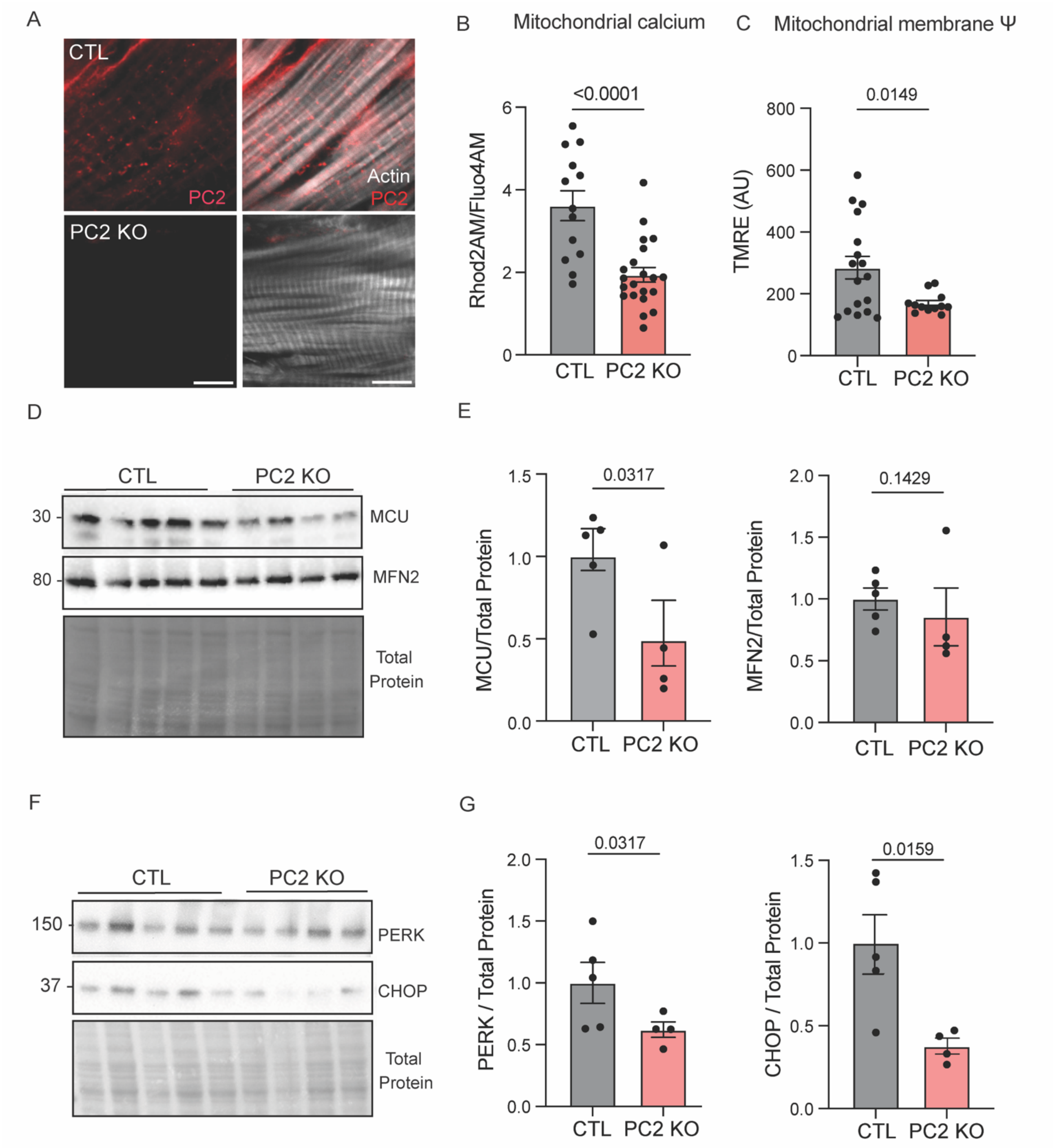


**Supplementary Figure 3. A.** Representative IF images of PC2 (red) and actin (white) in left ventricular sections from CTL and PC2 KO mice. Scale bar (5 µm). **B.** Analysis of mitochondrial calcium in isolated CTL and PC2 KO cardiomyocytes measured with Rhod2AM and normalized to Fluo4AM. N=3 animals per group, each dot represents a cell. Statistical analysis was assessed by using Mann-Whitney test. **C.** Analysis of mitochondrial membrane potential ($\Psi$) in CTL and PC2 KO cardiomyocytes measured with TMRE. N=3 animals per group, each dot represents a cell. Statistical analysis was assessed by using Mann-Whitney test. **D**. Protein expression of MCU and MFN2 from CTL (n=5) and PC2 KO (n=4) mice. Total protein used as loading control**. E**. Quantification of D. Statistical analysis was performed using Mann-Whitney test. **F**. Protein expression of PERK, CHOP from CTL (n=5) and PC2 KO (n=4) mice. Total protein used as loading control**. G.** Quantification of PERK and CHOP from F. Statistical analysis was performed using Mann-Whitney test.

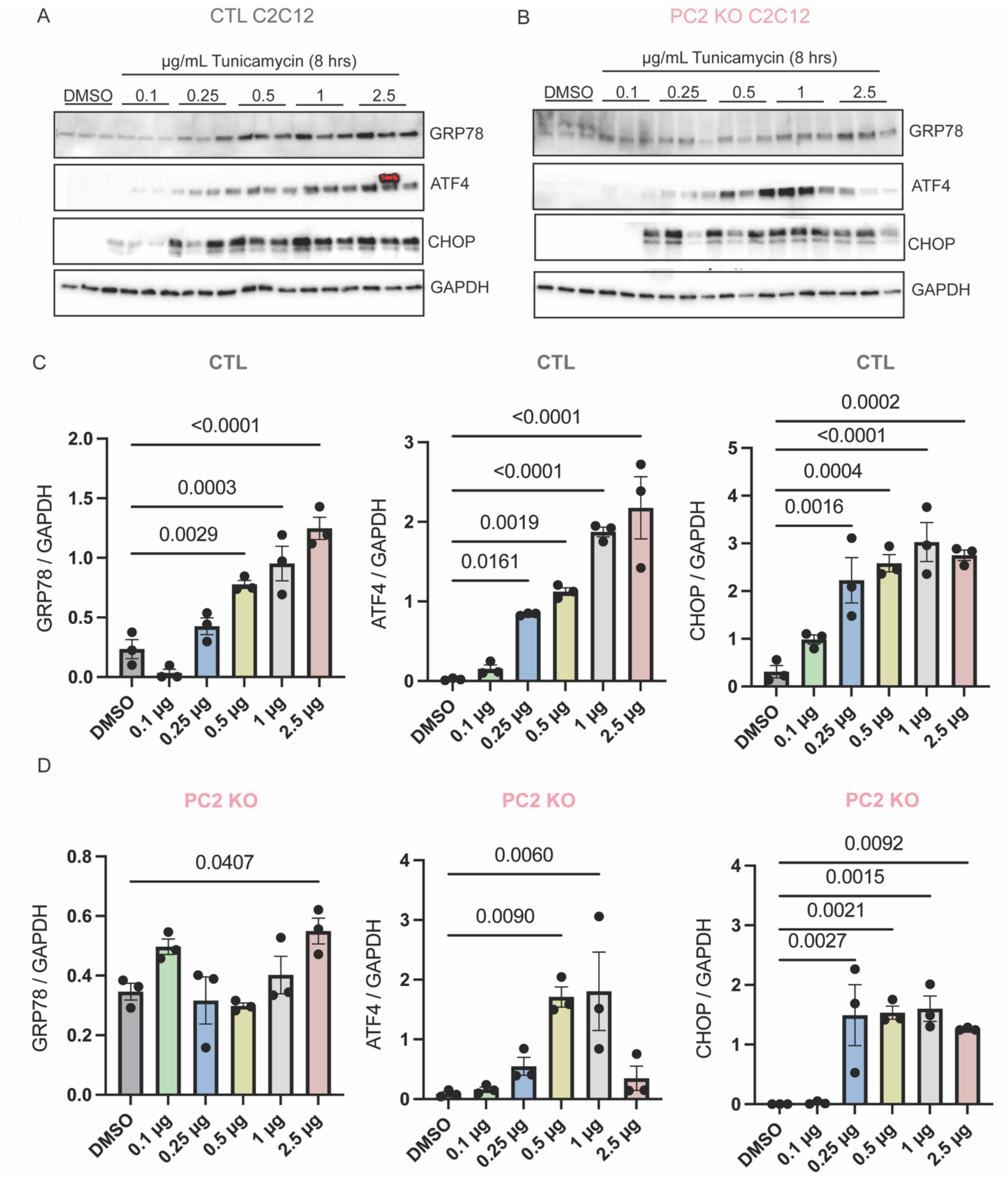


**Supplementary figure 4**: **A & C.** Protein expression and quantification of GRP78, ATF4 and CHOP expression in CTL C2C12 cells treated with increasing concentrations of TM (0.1, 0.25, 0.5, 1, and 2.5 µg/mL) for 8 hrs or DMSO (as a vehicle control). **B & D.** Protein expression and quantification of GRP78, ATF4 and CHOP expression in PC2 KO C2C12 cells treated with increasing concentrations of TM (0.1, 0.25, 0.5, 1, and 2.5 μg/mL) for 8 hrs or DMSO (as a vehicle control). Statistical analysis was assessed by using one-way ANOVA.

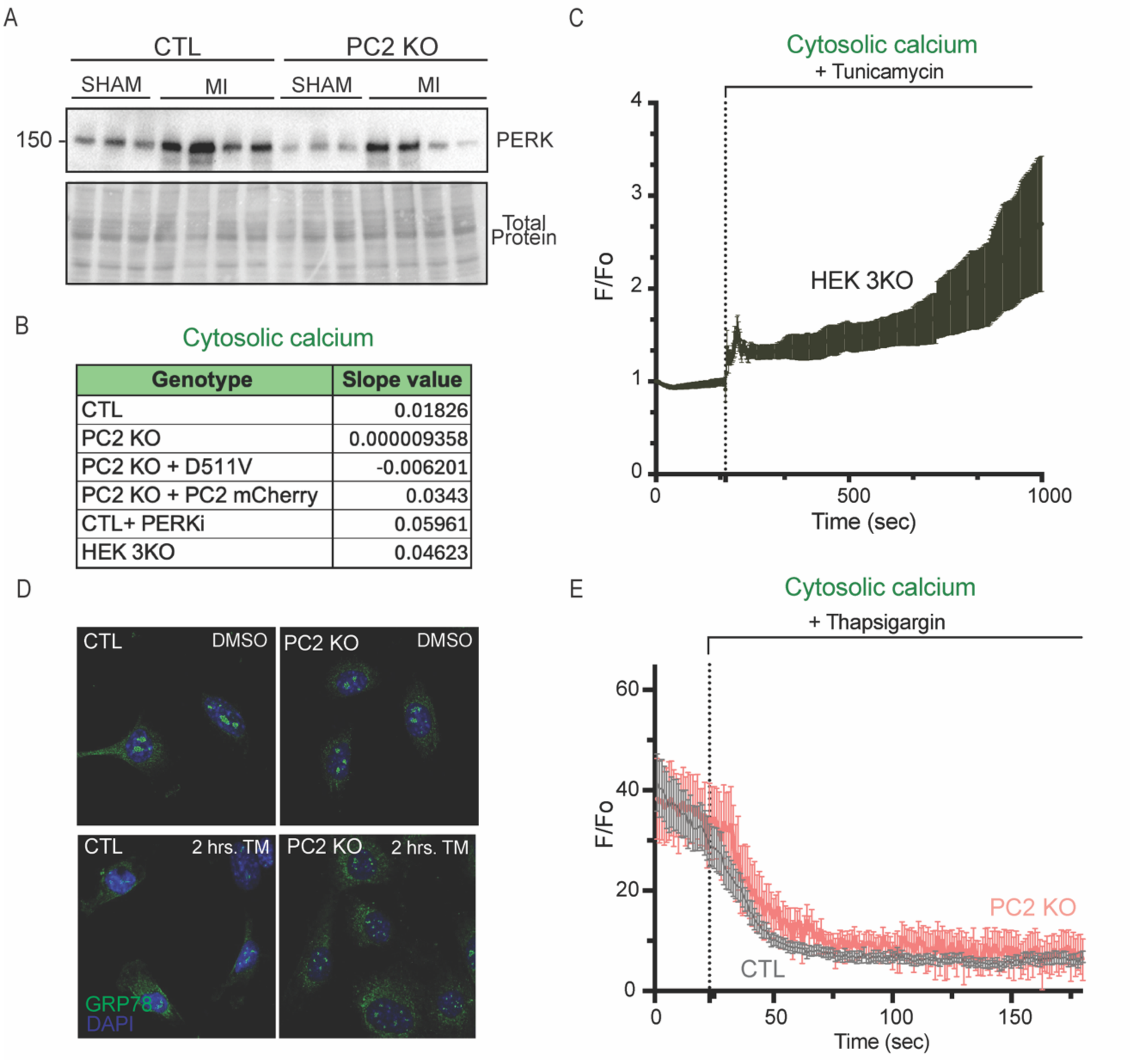


**Supplementary Figure 5**: **A**. Protein expression of PERK in CTL and PC2 KO SHAM (N=3) and MI (N=4) mice. **B.** Slope values from linear regression analysis of cytosolic calcium changes in CTL, PC2 KO, PC2 KO + D511V, PC2 KO + PC2 mCherry, CTL+ PERK inhibitor, and HEK3KO cells treated with TM (N=3). Analysis is fluorescence (y axis) divided by time in minutes (x axis). **C.** Averaged traces of cytosolic calcium (GCaMP6f) from HEK3KO cells treated with TM. **D.** Representative IF of GRP78 staining in CTL and PC2 KO cells treated with 2.5 μg/mL for 2 hours. **E.** Averaged traces of cytosolic calcium (GCaMP6f) from CTL and PC2 KO cells treated with 2 μM Thapsigargin. N=4.

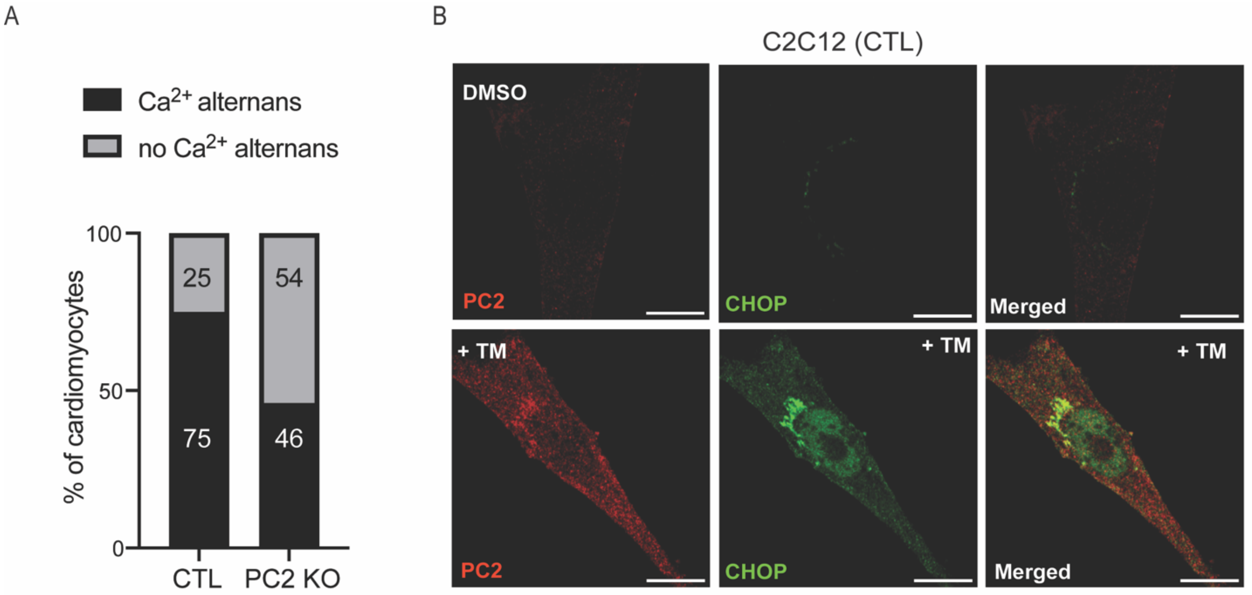


**Supplementary Figure 6: A.** Calcium alternans analysis in CTL and PC2 KO cells (17-25 cells per condition). N=3-5 animals per group. **B.** Representative IF images of PC2 and CHOP staining from CTL cells treated with DMSO or 2.5 μg/mL TM for 24hrs.
